# Predictive visual patterns during table tennis forehand rallies

**DOI:** 10.1101/2024.05.10.593485

**Authors:** Ryosuke Shinkai, Shintaro Ando, Yuki Nonaka, Tomohiro Kizuka, Seiji Ono

## Abstract

The purpose of this study was to clarify predictive visual patterns of skilled table tennis players during forehand rallies. Collegiate male table tennis players (n = 7) conducted forehand rallies at a constant tempo (100, 120 and 150 bpm) using a metronome. In each tempo condition, participants performed a total of 30 strokes (three conditions). Gaze fixation time, gaze targets and saccade eye movements were detected by video footage of an eye tracking device. We found that participants gazed at a ball approaching them only 20 % of the total rally time. Participants tended to gaze at the ball when the opponent hit the ball and move their gaze away from the ball after that. Furthermore, saccades were directed toward the opposite side of the court including the opponent after tracking the ball. These findings suggest that focusing on the opponent motion is important for successful forehand table tennis rallies. Taken together, skilled table tennis players are likely to use unique visual patterns for interceptive sports players to estimate spatiotemporal information about the ball.

## 1. Introduction

In racket-ball sports such as cricket (Mann et al., 2013), squash (Hayhoe, 2013), and tennis (Mann et al., 2019), skilled players direct their gaze to the future position of the approaching ball when it bounces. This is because the ball trajectory could suddenly change on bouncing.

Therefore, they attempt to anticipate the position where they can better judge the ball trajectory when it bounces. Although the time from bounce to hit has been reported as about 160, 240 and 320 ms (Mann et al., 2013), 400 to 800 ms (Hayhoe, 2013), and 300, 550 and 800 ms (Mann et al., 2019), the time in high pitch rallies in table tennis would be from 150 to 200 ms (Shinkai et al., 2023), meaning players would not afford to look at the ball in the time from bounce to hit. Therefore, the gaze pattern during table tennis rallies would differ from that of other racquet-ball sports based on several previous studies.

In table tennis, skilled players would not look at the ball just after the bounce because they could judge the ball trajectory earlier based on visual information about the hitting movement of an opposite player. Previous studies have suggested the importance of looking at the racket and swinging arm area of the opposite player for predicting the ball trajectories (Piras et al., 2016; Piras et al., 2019). Furthermore, Shinkai and colleagues (2022) have suggested that expert table tennis players took their gaze away from the ball earlier than semi-expert players during rallies, reflecting that skilled players are able to estimate the ball trajectory earlier than semi-skilled players. We defined this idiosyncratic gaze pattern during table tennis rallies as “predictive visual patterns,” although predictive visual patterns are generally interpreted as a gaze to the future position of a moving target (e.g., gaze position ahead of a ball approaching the player).

Thus, we attempted to expand the findings of previous studies by using constant forehand rallies in different tempo conditions.

Examining the saccade during intercept performance could be beneficial for an assessment of attention that cannot be assessed by fixation alone (Hoffman and Subramaniam, 1995).

Previous studies of interceptive sports have demonstrated that participants tracked a ball approaching them for as long as possible by predictive saccades, indicating that they attempted to predict the future location of the ball and acquire spatiotemporal information about the ball until the bat or racket hit the ball (Diaz et al., 2013; Mann et al., 2013; Mann et al., 2019).

Aoyama and colleagues (2022) have demonstrated that predictive saccades toward a moving target improve visuomotor performance according to the results from their original psychophysical experiment. These studies have suggested that the predictive saccades to the ball are effective for interception of the target. On the other hand, skilled table tennis players would make saccades toward their opponent during forehand rallies, rather than toward the ball approaching players. Thus, saccade analysis would explain where table tennis players direct their attention during the rallies.

The purpose of this study was to clarify predictive visual patterns of skilled table tennis players during forehand rallies. We attempted to quantify fixation, detailed gaze targeting a specific area (gaze target), and saccades during rallies. We raised the following three research questions (1) whether and, if yes, when do skilled table tennis players look at the ball when performing rallies, (2) whether the gaze target is only on the racket, swinging arm of the opposite player or location which the participant aims to hit the ball on the court, (3) whether they make saccades towards the opposite player rather than to where the ball will be after the bounce.

## 2. Materials and Methods

### 2.1. Participants

The participants were seven male college students who have participated in the All-Japan tournament (mean age: 19.7 ± 0.9 years, height: 169.9 ± 5.3 cm, body mass: 61.1 ± 4.2 kg, table tennis experience: 12.1 ± 2.4 years) and they reported having normal or corrected to normal vision and no known motor deficits. They were neither diagnosed with a stereoscopic problem nor strabismus. All participants gave their informed consent to participate in the experiment. This study was conducted in accordance with the 2013 Declaration of Helsinki, and all experimental protocols were approved by Research Ethics Committee at the Faculty of Health and Sport Sciences, University of Tsukuba. Written informed consent was obtained from all participants before their participation.

### 2.2. Experimental procedure

Participants wore an eye-tracking device (Pupil Invisible glasses, Pupil Labs, Berlin), and the calibration was performed before the experiment. In the calibration process, participants fixated the corners of the table as calibration grid to verify accurate tracking. We confirmed that the error in the gaze relative to each corner of the table was less than 1° of visual angle. The experimenter (Fig. 1-➀) conducted the forehand rallies as experimental tasks with participants (Fig. 1-➁). The experimenter delivered a ball to one target (diameter: 24cm, Fig 1-➂) drawn on the table court of the participant’s side, and the participant aimed to hit the ball to the circular target (diameter: 24cm, Fig. 1-➃) on the experimenter’s side. Participants conducted two trials to become familiar with this task. A metronome speaker (Creative MUVO 2c, CREATIVE, Japan, Fig. 1-➄) was set on the table near the net to accurately control the timing of each stroke. The three tempo conditions were conducted in the order of 100, 120 and 150 bpm. In each tempo condition, participants did 30 strokes (hitting of the ball) with the experimenter (3 conditions). In addition, four of seven participants started with the slowest tempo condition and proceeded to the faster conditions, while the remaining three participants started with the fastest condition and proceeded to the slower conditions.

**Fig. 1.**
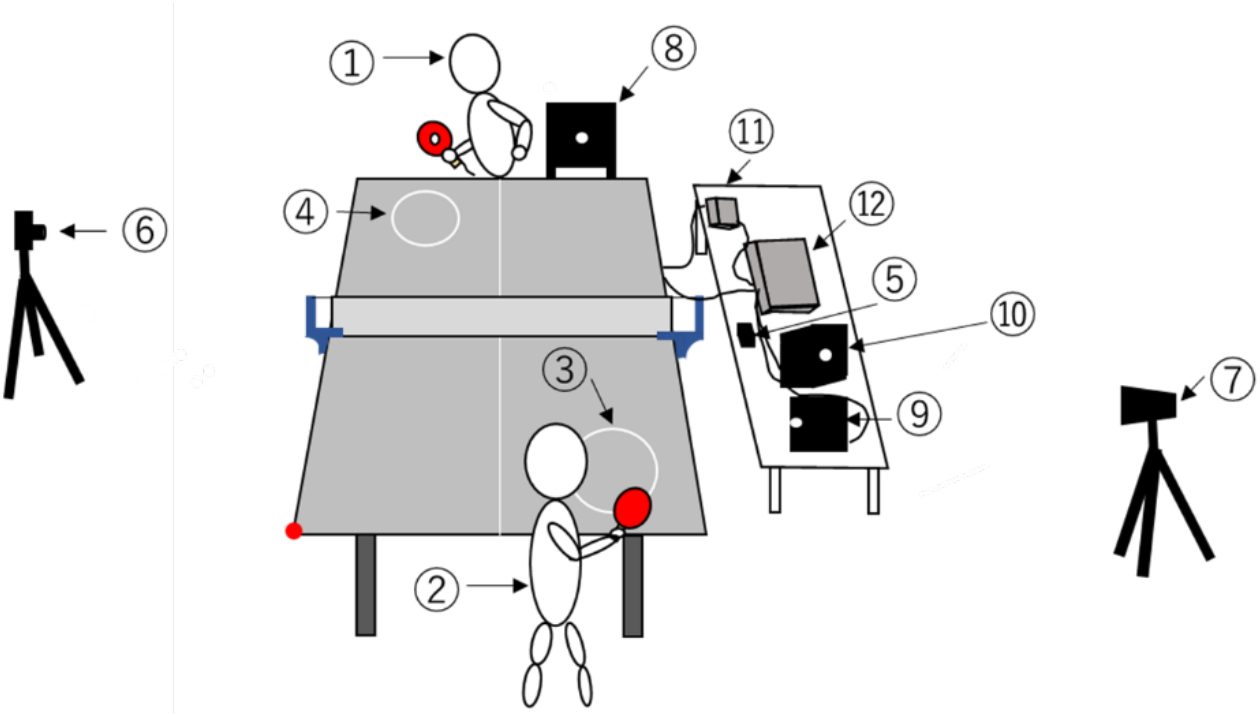
Overview of the experimental task. ➀skilled experimenter, ➁participant, ➂participant side of circular target, ➃experimenter side of circular target, ➄speaker, ➅➆high speed camera, ➇➈➉LED light, ⑪control box for gyro sensor, ⑫waveform generator. The red circular point at the corner of the participant side of the court shows the origin of the coordinate system to analyze ball trajectories during rallies.

### 2.3. Apparatus

An eye-tracking device was used for recording gaze targets of participants during experimental tasks. The images from the scene camera of the eye-tracking device were recorded at a sampling frequency of 30 Hz, and the eye movements the eye cameras were recorded at a sampling frequency of 200 Hz. Two high-speed cameras (frame rate =240 fps, EX-ZR200, CASIO, Japan, Fig. 1-➅, ➆) were set at the sides of the court for capturing ball trajectories during experimental tasks.

To synchronize the hitting time of the scene camera of the eye-tracking device and high-speed cameras, an acceleration sensor was attached to the rear of the racket of the experimenter, and the LED lights (Fig. 1-➇, ➈, ➉) were set to provide the signal output from the acceleration sensor (MP-A0-01A, MicroStone, Japan, Fig. 1-⑪) through the waveform generator (SG-4211, IWATU, Japan, Fig. 1-⑫). Therefore, the LED lights flashed with the vibration at the moment the experimenter hit the ball. The delay from the hitting time to the LED flash was < 5 ms. The time when the LED lights flashed was captured from each image of the high-speed camera (frame rate =240 fps, EX-ZR200, CASIO, Japan, Fig. 1-➅, ➆). The images from the scene camera of the eye-tracking device were recorded at a sampling frequency of 30 Hz, and the eye movements were recorded at a sampling frequency of 200 Hz.

### 2.4. Data analysis

Video footage from the scene camera of the eye-tracking device was digitized using motion analysis software (Frame-DIAS IV, DKH, Japan) to determine the coordinates (pixels) of a ball position relative to the head in each frame of the video footage. Ball coordinates were resampled from 30 to 200 Hz by using second or third-order spline interpolation to match them with eye direction data. The pixel values of the coordinates were converted to visual angles based on the specifications of the eye-tracking device (horizontal angle: 82 deg / 1088 px, vertical angle: 82 deg / 1080 px). The coordinate origin of the scene camera was defined as the center screen of video footage.

To evaluate detailed gaze targets during rallies, the visual search behavior was analyzed for individual participants, consisting of the following analyses:

#### Number of Fixations per trial

The mean number of fixations per trial was calculated. A fixation was defined as gaze being maintained at a single area of interest, either stationary or moving, for a minimum of 120 ms (van Biemen et al., 2022) with a visual angle of less than 3° (Rodrigues et al., 2002; Piras et al., 2016). Although this study did not provide a speed criterion for the detection of fixation, gazing at a single area of interest within the same 3 deg, or for at least 120 ms reflects that gaze clearly stays on the target. Figure 2 shows raw data of eye and ball position in the head coordinate system when the fixation was detected. A moving fixation can be considered as a smooth pursuit to follow a moving ball (Lisberger et al., 1987).

**Fig. 2.**
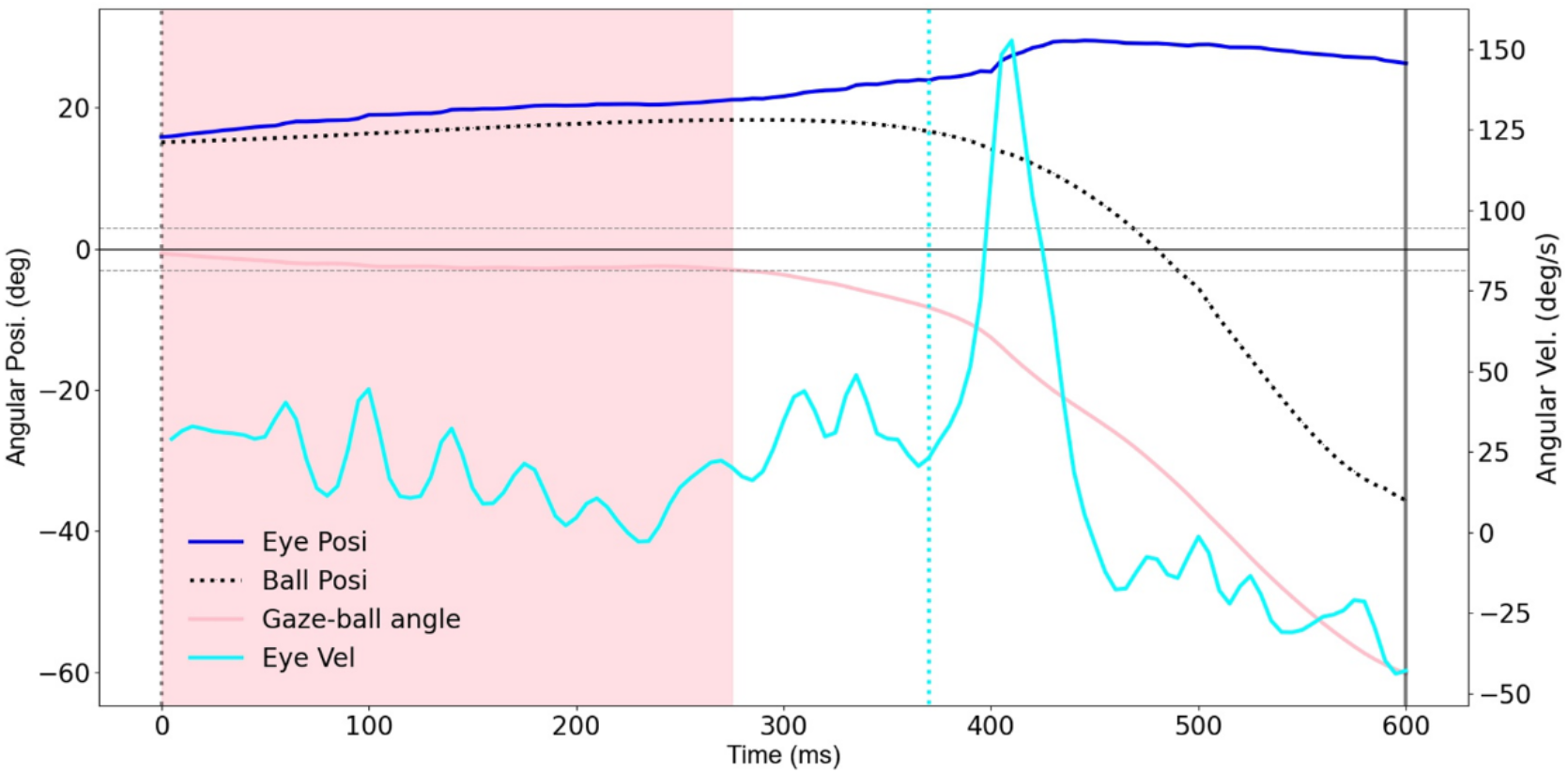
Typical waveforms of eye, ball and gaze-ball angle in the 100 bpm condition. A dotted vertical line along 0 ms indicates experimenter’s hit. The solid vertical line along the latest time indicates participant’s hit. A pink area indicates fixation duration. Horizontal dotted lines indicate ± 3 degree of visual angle for detection of the fixation offset. A lightblue dotted line indicates the onset of a saccade. Upward deflections of eye and ball traces indicate leftward movements.

#### The Fixation time to the ball approaching participants

The relative time (%) of fixation time in ball approach phase were calculated based on the above definitions.

#### Gaze targets

Eight areas of interest where gaze could be directed were defined: the experimenter’s racket, the ball, the opposite court near the experimenter (Near), the opposite court on the middle line (Middle), the opposite court far from the experimenter (Far), the circular target on the opposite court, the space between circular target and racket and the trunk of experimenter (Fig. 3). The reason which we divided the table into three parts was to clarify whether gaze directions were close to the ball or opponent. All gaze targets were determined whether the difference between the coordinate of the gaze target and a certain point was within 3° of the visual angles. If the ball overlapped the position of the racket when the experimenter hit the ball, we assigned the gaze to the position of the ball. If the ball overlapped the position of the target, the gaze was assigned to the position of the ball. Furthermore, if the racket overlapped the position of the experimenter, especially for the body trunk, we assigned the gaze to the position of the racket. However, our hypothesis was that the gaze pattern would not look at the ball. Thus, this choice is conservative because it is against our preferred interpretation. Data analysis of gaze targets was performed by manually encoding frame-by-frame video images from the eye-tracking device. After collecting all data, each gaze target in the relative time (0 – 100 %) was determined by statistical mode. In other words, we determined the most common region to be fixated at each percentage of time for each participant, and then determined the percentage for each region across participants. This process allows us to indicate detailed gaze targets during rallies, although the gaze targets do not necessarily mean “fixation”. It also allows comparison of gaze targets among all participants.

**Fig. 3.**
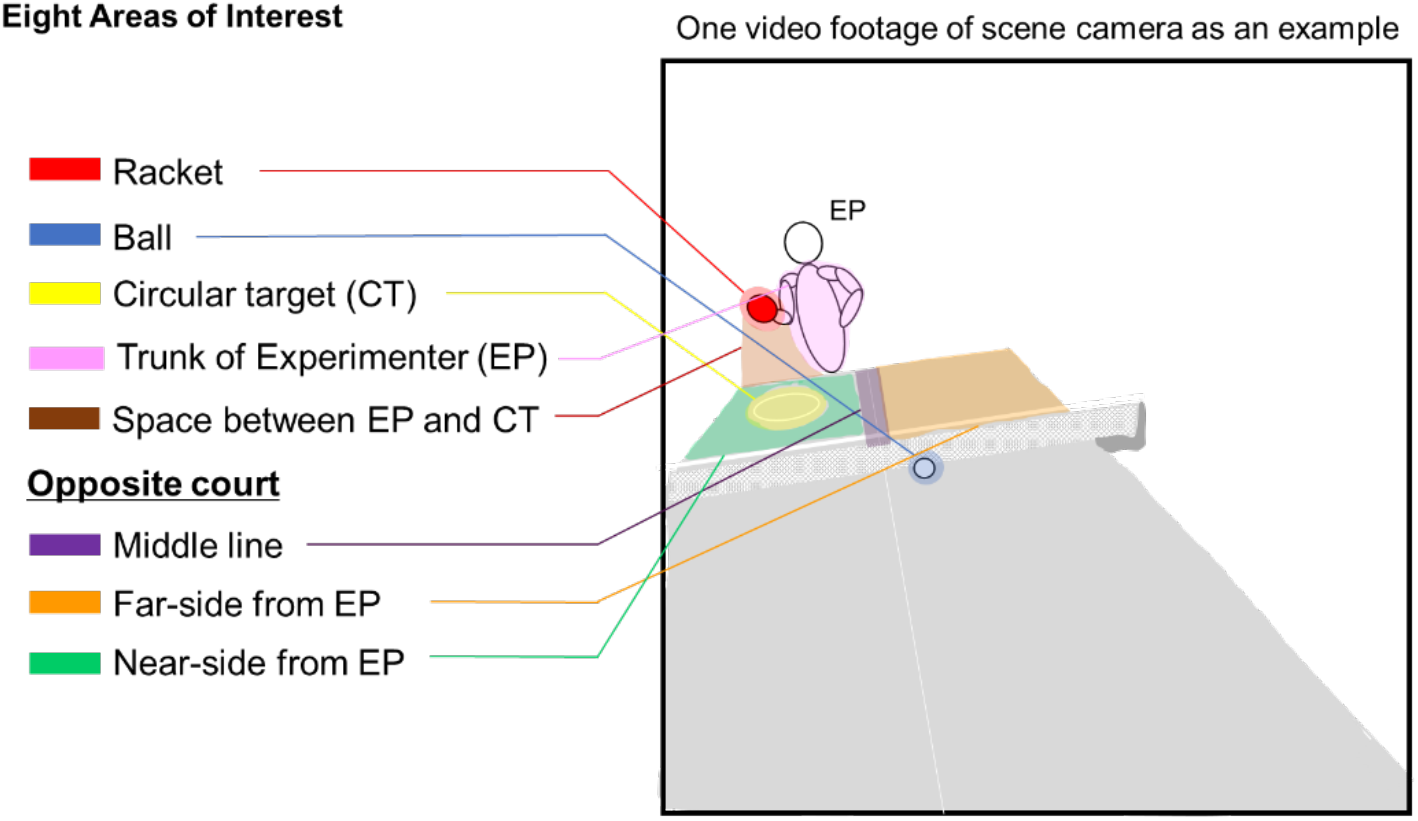
Eight defined areas of interest in this study. Eight areas of interest were defined: the racket, the ball, the opposite court near the experimenter (Near), the opposite court on the middle line (Middle), the opposite court far from the experimenter (Far), the circular target on the opposite court, the space between circular target and racket and the trunk of the experimenter.

To evaluate the saccades in each shot, the distance between the eye direction on the next and previous samples was divided by the time between those samples to determine eye velocity at each moment. We detected saccades in horizontal and vertical directions because the eye position data was exported separately in the horizontal and vertical direction. We defined the onset of saccades as the portion of 5 % of the peak value of saccadic eye velocity. Peak eye velocity during saccades should be greater than 100 deg/s. Moreover, the amplitude of eye direction during saccades should be at least 1 degree to avoid the detection of smooth pursuit (Mann et al., 2019). All gaze targets immediately after saccades showed one of the eight defined areas of interest. Therefore, the gaze targets immediately after saccades were categorized into eight areas of interest.

Fixation, gaze targets and saccades were defined in the head coordinate system. Although we captured head movements of participants with a build-in gyroscope of the eye-tracking device (sampling rate: 200 Hz), only a small amount of head rotation was detected (horizontal average angle: -4.7 ± 2.7 degrees, vertical average angle; -1.6 ± 2.2 degrees). In this case, negative values indicate a rightward deflection in the horizontal direction and a downward deflection in the vertical direction.

To evaluate whether the participant’s hits were successful in terms of the racket hitting the ball and of the ball landing at the target area, the images of two high-speed cameras (frame rate =240 fps, EX-ZR200, CASIO, Japan, Fig. 1-➅, ➆) were used for constructing three dimensional coordinates of ball trajectories. The images at the moment the ball was landing at the experimenter’s side of the court were analyzed to calculate the distance relative to the center of the circular target, reflecting the hitting accuracy of participants.

In this study, the data about the gaze targets and ball trajectories were presented relative to a normalized time. The normalized time begins at the moment when the experimenter hits the ball toward the participant (time = 0 %), and ends at the moment the experimenter hits the ball back again (time = 100 %), after the participant has returned it to him (time = 50 %). The normalized time was implemented by the high-speed camera (Fig. 1-➅). The onset of the experimenter’s hit was inferred from the LEDs on his racket whereas the onset of the participant’s hit was inferred from the images of the high-speed camera, capturing not only ball trajectories but also the racket movements of the experimenter and participants.

### 2.5. Statistical analysis

To examine a difference in relative fixation time between tempo conditions, a one-way ANOVA was performed. To compare the percentage of gaze targets on each area of interest across normalized time, a one-way ANOVA was performed. The ANOVA reflects where the gaze targets toward each area of interest occurred significantly at what percentage of the normalized time. Post hoc comparisons were Bonferroni adjusted to test all combinations across gaze target data at each normalized time bin. An alpha level of 5 % was applied for all the tests. Effect sizes were calculated as partial eta squares. All statistical tests were conducted by IBM SPSS software version 27 (SPSS Inc, USA). The obtained valuables were calculated separately for each trial and then averaged across trials of each condition.

### 2.6. Properties of ball trajectories hit by the experimenter

Two components were analyzed, (1) time from one player’s hitting to another player’s hitting during rallies and (2) ball positions relative to the circular target to confirm the reliability of the test. For the ball trajectory, the position when the ball bounced in all trials was calculated by using motion analysis software (Frame-DIAS IV, DKH, Japan). Two high-speed cameras (frame rate = 240 fps, EX-ZR200, CASIO, Japan, Fig. 1-➅, ➆) were used for capturing ball trajectories. The origin of the coordinate system was the corner of the participant side of the court (red point in Figure 1). The time from the experimenter’s hitting to the participant’s hitting was 576 ± 15.6 ms in the 100 bpm, 486 ± 13.8 ms in the 120 bpm and 391 ± 10.7 ms in the 150 bpm. The time from the participant’s hitting to the experimenter’s hitting was 617 ± 14.0 ms in the 100 bpm, 507 ± 12.3 ms in the 120 bpm and 401 ± 11.0 ms in the 150 bpm. These results indicate that forehand rally tasks were successfully conducted in each condition.

All trials across participants were successful in terms of the racket hitting the ball. On the other hand, not all hits were successful in landing on the circular target. The average number of successful balls landing on the circular target in all tempo conditions was 16.4 ± 4.6 hits in the 100 bpm, 17 ± 3.7 hits in the 120 bpm and 12 ± 4.6 hits in the 150 bpm, respectively. However, figure 4 indicates that plots of ball positions on the court of the experimenter’s side are scattered very close to the circular target or with a small area within the circular target. Furthermore, we examined whether the successful hitting was related to gaze deployment. Figure 5 indicated that gaze deployment on the hit was not related to that on misses.

**Fig. 4.**
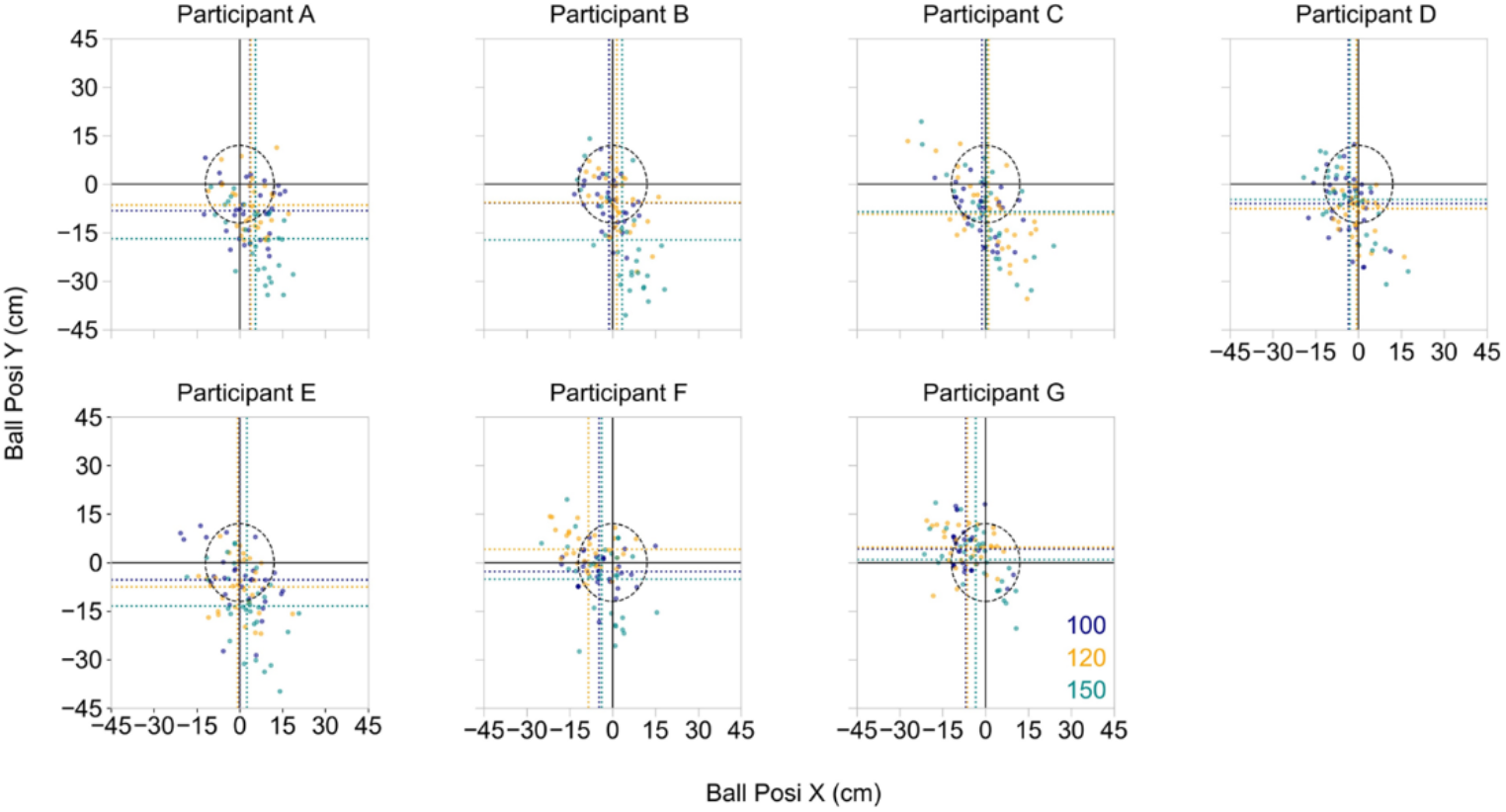
Ball positions relative to the center coordinate of the circular target at each participant when the ball bounced on the experimenter’s court. Darkblue, orange and darkcyan dots indicate single trial data in the 100, 120 and 150 bpm, respectively. The horizontal and vertical dotted lines indicate the mean values of the ball position at each participant.

**Fig. 5.**
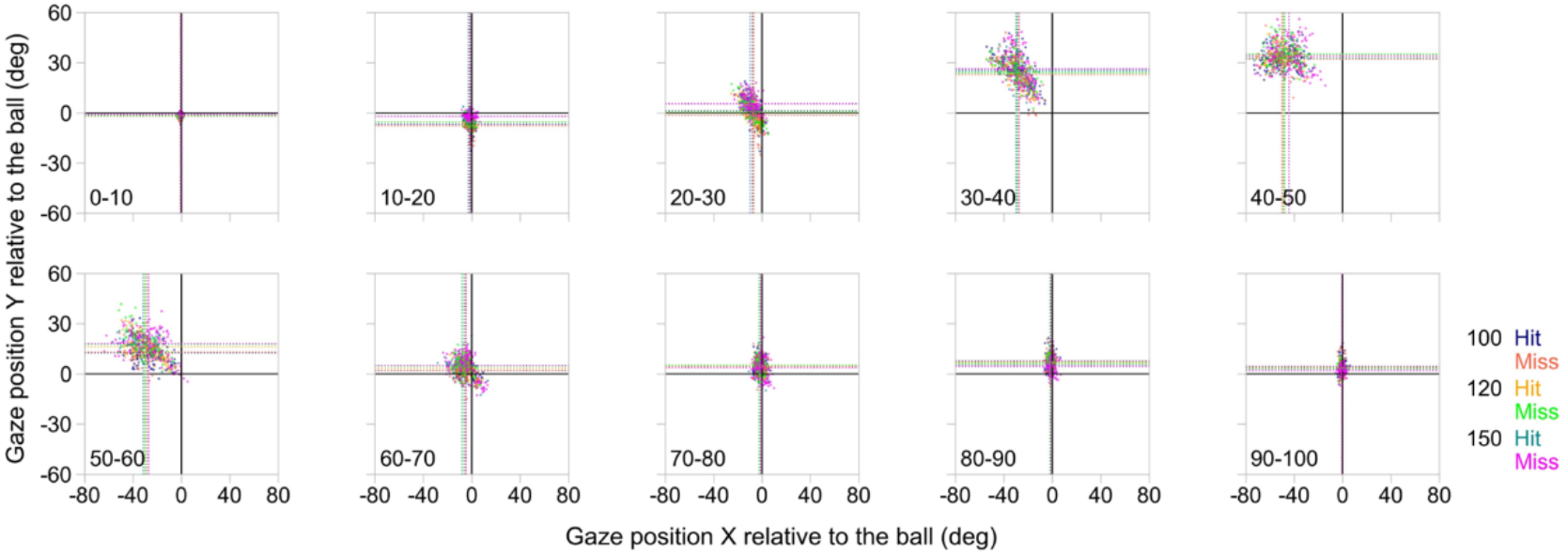
Gaze positions relative to the ball position in each normalized time point. Darkblue, orange and darkcyan dots indicate single trial data for “Hit” whereas salmon red, lime green and magenta dots indicate single trial data for “Miss” in the 100, 120 and 150 bpm, respectively. The horizontal and vertical dotted lines indicate the mean values of the ball position at each participant.

## 3. Results

### 3.1. Occurrence of fixations on the ball during rallies

Fixations on the ball occurred in two phases around the hitting of the experimenter. The first phase is the time period immediately after the experimenter hit the ball, meaning that the ball was approaching the participant. The second phase is the time period immediately before the ball arrived at the experimenter, meaning the period from the time the participant hit the ball until the experimenter hit the ball back.

During the ball approach phase, participants performed on average 21.1 ± 9.0 fixations in the 100 bpm (total: 148), 19.0 ± 10.1 fixations in the 120 bpm (total: 133) and 14.6 ± 9.1 fixations in the 150 bpm (total: 133). The rate of fixation to the ball approaching participants was 70.5 % in the 100 bpm condition, 63.3 % in the 120 bpm condition and 47.6 % in the 150 bpm condition. These results indicate that the occurrence of fixations on the ball approaching participants decreased as the tempo was increased.

Immediately before the experimenter hit the ball, participants performed on average 12.7 ± 5.2 fixations in the 100 bpm (total: 89), 14.1 ± 8.2 fixations in the 120 bpm (total: 99) and 14.0 ± 5.1 fixations in the 150 bpm (total: 98). The rate of fixation on the ball approaching the experimenter was 42.4 % in the 100 bpm condition, 47.1 % in the 120 bpm condition and 46.7 % in the 150 bpm condition. These results do not indicate that the occurrence of fixations on the ball immediately before the experimenter hit the ball changed as the tempo increased or decreased.

### 3.2. Fixation time

The mean relative fixation time on the ball approaching participants in each tempo condition was 19.6 ± 2.4 % in the 100 bpm condition, 19.5 ± 1.5 % in the 120 bpm, and 19.4 ± 1.1 % in the 150 bpm (Fig. 6A-6C). There was no significant main effect of the tempo condition on the relative fixation time. The results indicate that relative fixation time was not significantly decreased as the tempo increased.

**Fig. 6.**
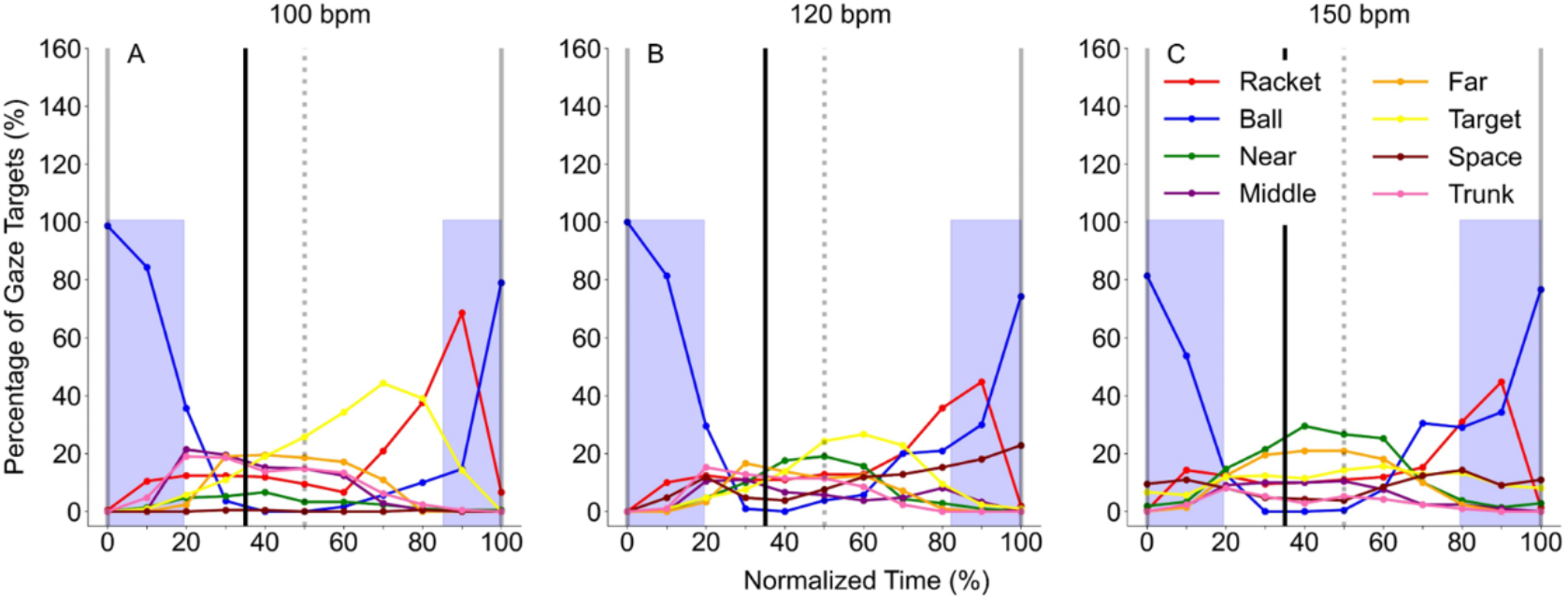
Average percentage of gaze targets at each normalized time in all tempo condition. The X-axis shows normalized time from 0 to 100 %. The normalized time begins at the moment when the experimenter hits the ball toward the participant (time = 0 %), and ends at the moment the experimenter hits the ball back again (time = 100 %), after the participant has returned it to him (time = 50 %). The black dotted vertical line along 35 % of normalized time indicates the moment at which the ball bounced on the experimenter’s side of the court. The Y-axis shows the percentage of gaze target at each normalized time. Blue shaded areas show averaged fixation durations on the ball.

The mean relative fixation time on the ball approaching the experimenter in each tempo condition was 14.7 ± 4.6 % in the 100 bpm condition, 17.8 ± 5.5 % in the 120 bpm, and 20.5 ± 4.9 % in the 150 bpm (Fig.6A-6C). There was no significant main effect of the tempo condition on the relative fixation time. These results indicate that relative fixation time was not significantly decreased as the tempo increased.

### 3.3. Gaze targets during rallies

Gaze targets during forehand rallies in this study demonstrate the defined eight areas of interest. After fixation on the ball approaching participants, most of the gaze targets stayed on the opposite court in each tempo condition (100bpm; 99 % of total trials, 120bpm; 100 % of total trials, 150 bpm; 97.7 % of total trials). Figure 6 indicates an example of gaze behavior during rallies. The point of gaze stayed at the participant’s side of the court in all video frames, indicating that the participant looked away from the ball approaching him sooner after the experimenter hit the ball.

A one-way ANOVA in the 100 bpm condition showed that the percentage of gaze target to the ball at each time in the 20 to 90 % was significantly lower than that at the time of 0 and 100 % (*p* < 0.01, Fig.6A). A one-way ANOVA in the 120 bpm condition showed that the percentage of gaze target for the ball at each time in the 20 to 80 % was significantly lower than that at the time of 0 and 100 % (*p* < 0.01, Fig.6B). A one-way ANOVA in the 150 bpm condition showed that the percentage of gaze target for the ball at each time in the 30 to 60 % was significantly lower than that at the time of 0 and 100 % (*p* < 0.01, Fig.6C). Overall, these results indicate that participants direct their gaze to the ball nearly around the time that the experimenter hit the ball and gradually shift their gaze to other defined areas of interest. However, there was no significant difference in the percentage of gaze target to other defined areas of interest time-by-time in any of the tempo conditions. This result indicates that the gaze targets after moving away from the ball approaching participants varied among individual participants, even though the gaze is directed to the opposite side of the court during rallies. Figures 7, 8 and 9 show the gaze targets at each normalized time per trial in all tempo conditions. These figures, indeed, indicate that gaze targets after moving away from the ball approaching participants varied among individual participants.

**Fig. 7.**
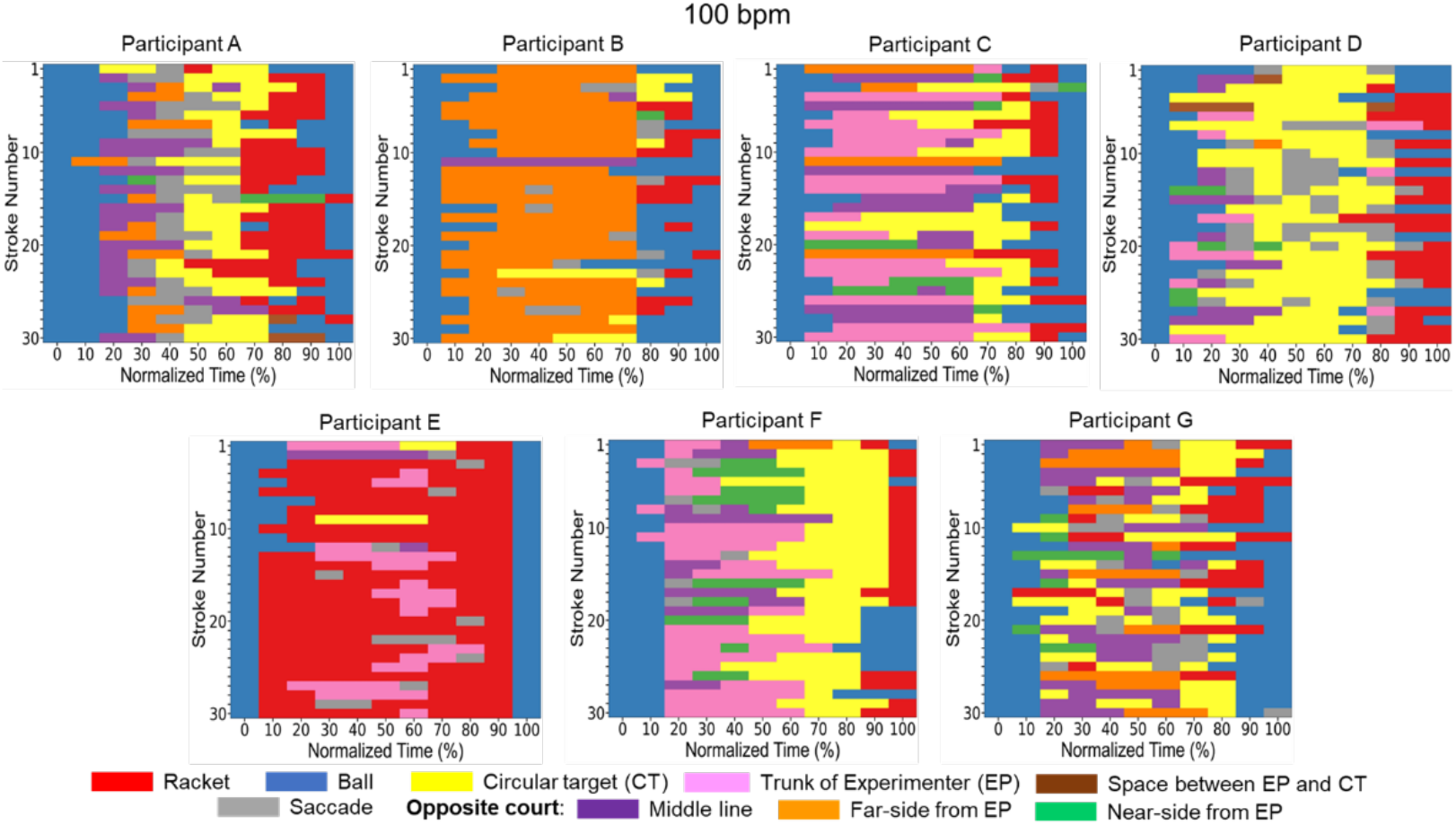
Individual gaze directions at each normalized time in the 100 bpm condition. The definition of the X-axis is the same as in Fig. 6. The Y-axis shows the stroke number from 1 to 30 strokes.

**Fig. 8.**
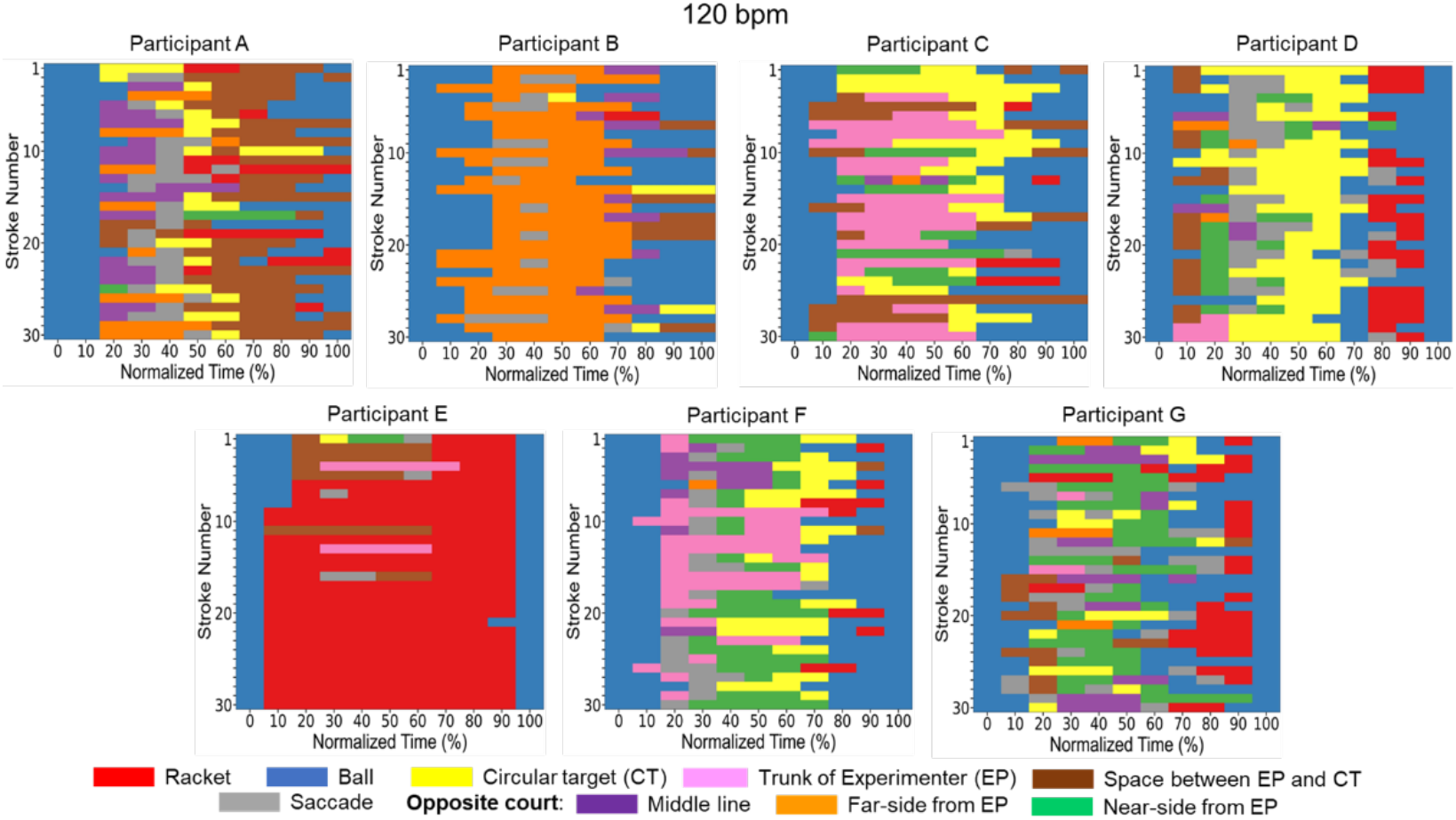
Individual gaze directions at each normalized time in the 120 bpm condition. The definition of the X- and Y-axis are the same as in Fig. 7.

**Fig. 9.**
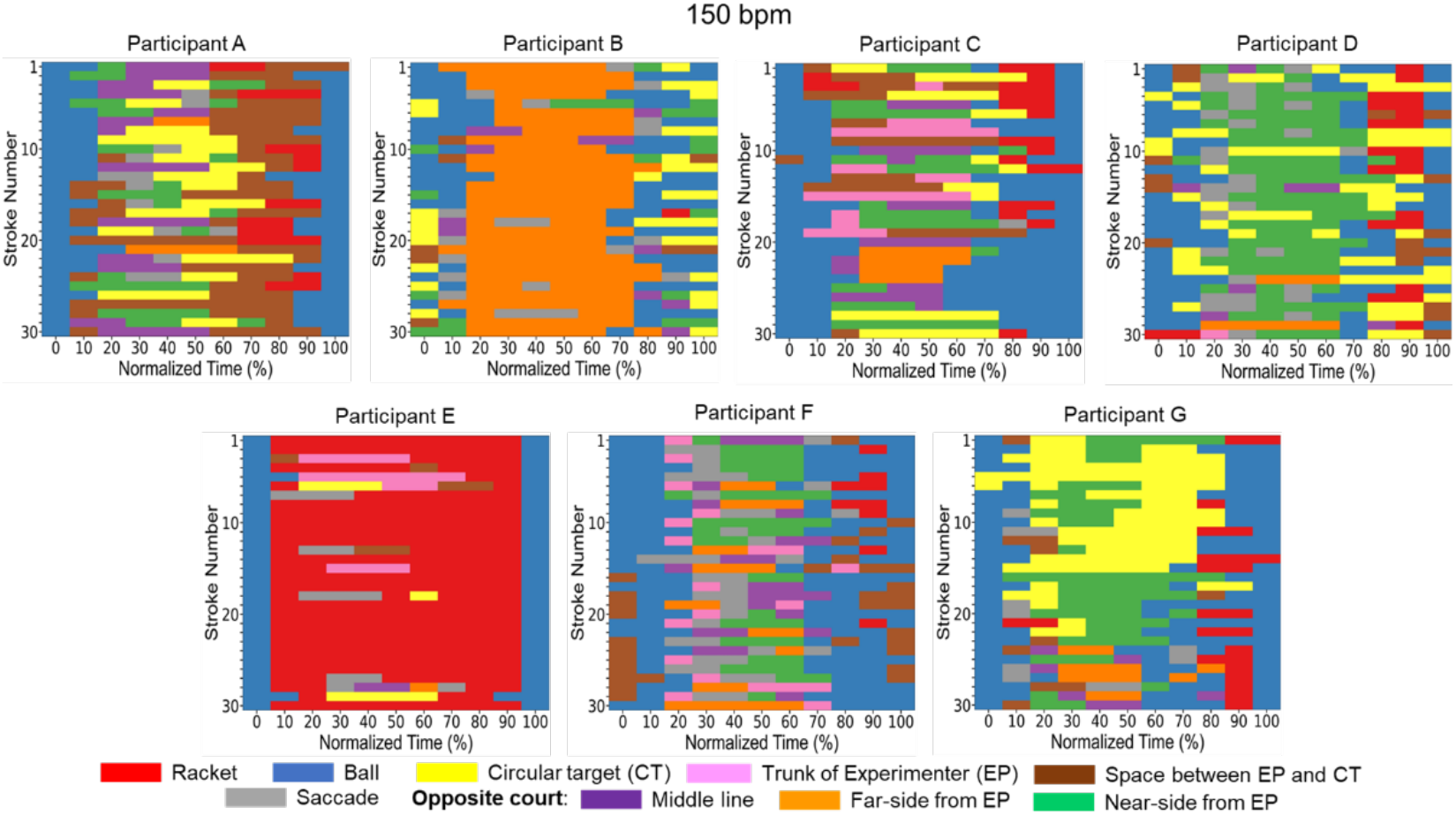
Individual gaze directions at each normalized time in the 150 bpm condition. The definition of the X- and Y-axis are the same as in Fig. 7.

### 3.4. Gaze targets immediately after saccades

Table 1 shows the number of horizontal and vertical saccades immediately before gazing at each defined area of interest in all participants. In the horizontal direction, participants performed on average 57.6 ± 25.2 saccades in the 100 bpm (total: 387), 57.7 ± 30.2 saccades in the 120 bpm (total: 376) and 53.3 ± 32.4 saccades in the 150 bpm (total: 385). On the other hand, In the vertical direction, participants performed on average 40.6 ± 15.8 saccades in the 100 bpm (total: 282), 47.0 ± 24.9 saccades in the 120 bpm (total: 252) and 29.9 ± 19.7 saccades in the 150 bpm (total: 207). These results indicate that the occurrence of saccades for the ball is not frequent, suggesting most of the attention in rallies focuses on the opposite direction even in the ball approach phase. Figure 10 shows the occurrence of saccades for each defined direction in each normalized time. The gaze targets after horizontal and vertical saccades in the ball approach phase (in the time of 0 -50 %) tended to be other defined areas of interest away from the ball, suggesting that not only the gaze targets but also the attentional directions were opposite side of the court.

**Table 1.** The number of saccades in horizontal and vertical directions immediately before gazing in each direction.

|  |  | Racket | Ball | Near | Middle | Far | Target | Space | Trunk |
| --- | --- | --- | --- | --- | --- | --- | --- | --- | --- |
| 100bpm | Horizontal | 70 | 33 | 42 | 35 | 49 | 95 | 43 | 20 |
|  | Vertical | 64 | 44 | 40 | 8 | 24 | 59 | 30 | 13 |
| 120bpm | Horizontal | 59 | 18 | 70 | 30 | 47 | 90 | 44 | 18 |
|  | Vertical | 101 | 34 | 29 | 27 | 3 | 31 | 27 | 0 |
| 150bpm | Horizontal | 65 | 33 | 79 | 33 | 74 | 68 | 28 | 5 |
|  | Vertical | 22 | 10 | 81 | 13 | 39 | 26 | 13 | 3 |

**Fig. 10.**
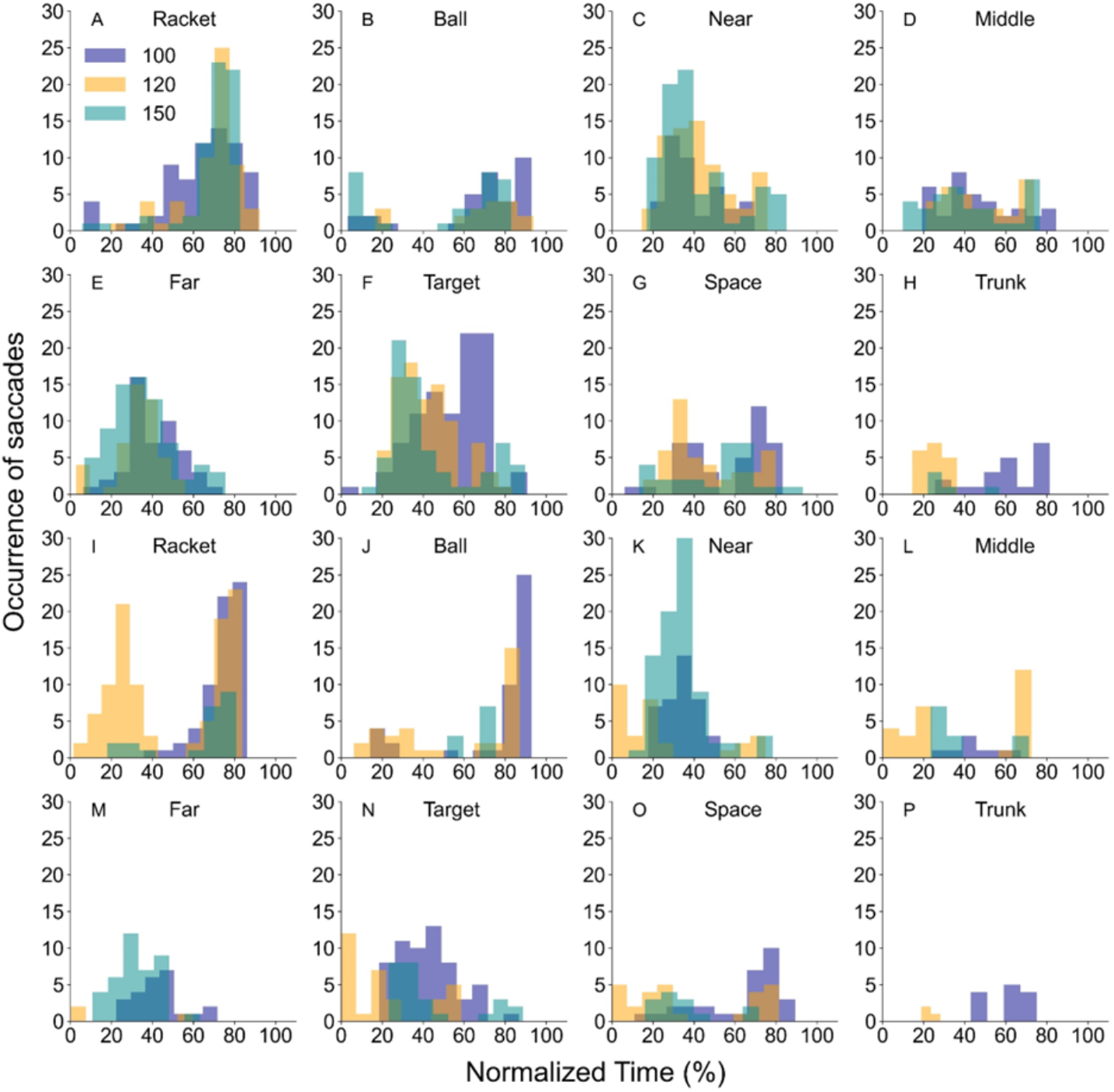
Occurrence of horizontal saccades (Fig. 10A-10H) and vertical saccades (Fig. 10I-10P) for each defined gaze target in each normalized time in all tempo condition.

### 3.5. Relationship between hitting accuracy and gaze deployment

All trials across participants were successful in terms of the racket hitting the ball. On the other hand, not all hits were successful in landing on the circular target. The average number of successful balls landing on the circular target in all tempo conditions was 16.4 ± 4.6 hits in the 100 bpm, 17 ± 3.7 hits in the 120 bpm and 12 ± 4.6 hits in the 150 bpm, respectively. However, figure 4 indicates that plots of ball positions on the court of the experimenter’s side are scattered with a small range very near or within the circular target. Furthermore, we also examined whether the successful hitting was related to gaze deployment. Figure 5 indicates that gaze deployment for the hit was not related to that for miss.

## 4. Discussion

In this study, we examined the predictive visual patterns of skilled table tennis players while they conducted forehand rally tasks. We found that participants did fixate on a ball approaching participants only 20 % of the total rally time. In addition, the time period in which they gazed at the ball during rallies was when the opponent hit the ball. This result reflects that they look away from the ball approaching them. However, there was no difference in gaze target other than the ball, indicating that gaze target after fixation on the ball was not consistent among individual participants, although their gaze was directed to the opposite side of the court during rallies. Furthermore, the occurrence of the saccade for the ball is not frequent, suggesting most of the attention during rallies was focused in the opposite direction. Taken together, skilled table tennis players move their gaze away from the ball in the earlier phase of ball approach, which could be due to predictive visual patterns during rallies.

### 4.1. Fixation on the ball during rallies

Fixation on the ball was observed immediately before and after the experimenter hit the ball in all tempo conditions. Furthermore, the relative ball fixation time was only about 20 % of the total rally time in all tempo conditions. These results support that it is important for table tennis players to gaze at the ball in the initial ball approach phase (Ripoll and Fleurance, 1988, Shinkai et al., 2023). Rodrigues and colleagues (2002) have also suggested that participants move their gaze away from the ball in the earlier phase of ball approach, supporting our results in this study. They conducted an experiment in which participants returned the ball to the right or left target cue under three different timing cue conditions (pre-, early- and late-cue conditions). In their study, the cue light was illuminated before the serve (pre-cue), during the initial phase of the ball flight (early-cue) or during the last phase of the ball flight (late-cue). In particular, they have shown that the gaze-ball angles in the early-cue and late-cue are more than 3 degrees after 20 % of the trial time. In other words, participants in their study would have to judge which cue is lit by gazing at the opposite court until the cue was lit. Therefore, the gaze target would remain at the opposite court even though the ball was approaching the participants. This study is the first to show similar results with a constant forehand rally task.

### 4.2. Gaze targets during rallies

Most of the gaze targets during rallies pointed to the eight defined areas of interest, indicating that gaze targets of participants dwelled on the opposite side of the court, even in the ball approach phase. Participants directed their gaze to the ball immediately before and after the experimenter hit the ball and gradually shifted their gaze to other defined areas of interest. These results support the results of fixation on the ball as mentioned above. Furthermore, these results suggest where skilled table tennis players direct their gaze after fixation on the ball in the ball-tracing phase. Shinkai and colleagues (2022) have indicated that the gaze-ball angle of expert players is significantly larger than that of semi-expert players, suggesting that expert players looked away from the ball approaching them. However, it is uncertain where expert players gaze after fixation on the ball. Therefore, the present study provides valuable insights for further understanding of predictive visual patterns. In particular, these results are surprising because they did not move their gaze to where the ball bounced. Where athletes make the predictive saccade after the ball bounces has been discussed in previous studies (Diaz et al., 2013; Mann et al., 2013; Mann et al., 2019). These studies have indicated that predictive saccades are made to a point that the ball will reach after the ball bounces. However, the result in this study is different from previous studies. This discrepancy would come from the small size of a table tennis court, causing severe time constraints for table tennis players to hit the ball approaching them. Furthermore, the participants in this study had expert skills for the rally tasks since they afforded to execute successful rallies even while gazing at the opposite’s side of the court.

Previous studies for baseball (Nakamoto et al., 2022) and softball (Takamido et al., 2022) have indicated that skilled players estimate the ball speed based on kinematic information of the pitching motion. In table tennis, it is likely important to acquire kinematic information about the opponent to estimate not only the ball speed, but also the ball direction (e.g., right or left). In particular, Piras and colleagues (2016) have suggested that acquiring the kinematic information in the hand-racket area improves reaction time and accuracy to the direction of the ball hit by the opponent. Thus, it is suggested that acquiring the kinematic information of the opponent is related to predictive visual patterns during forehand table tennis rallies. Figures 7, 8 and 9 indicate where the gaze targets of each participant are at each normalized time. All participants tended to direct their gaze on the ball along 0, 10 and 100 % of normalized time. Although there are large individual differences in where participants looked, the certain finding is that participants looked at the ball approaching them shorter due to the prediction of where the ball will come. Large individual differences in gaze targets would be associated with differences in prediction on the ball during rallies. For example, as shown in figures 7, 8 and 9, gaze targets in participant B mostly indicate far-side from the experimenter after fixation on the approaching ball, reflecting that this participant could make use of a visual-pivot strategy to watch the whole visual scene, especially for the opponent area during rallies. The visual pivot strategy is one of the best strategies to use peripheral vision during the performance (Williams et al., 1999; Kato, 2020). On the other hand, gaze targets in participant G are scattered around the opponent area and are not consistent across tempo conditions. Although the gaze targets of this participant did not match with the circular target and racket-arm area of the experimenter, he would make use of peripheral information about them. Actually, his hitting performance was the best of all participants (Fig. 4).

### 4.3. Occurrence of saccades during rallies

Participants in this study often made saccades to gaze toward the other defined areas of interest (experimenter’s direction) before and after fixation on the ball. Hoffman and Subramaniam (1995) have suggested that visuospatial attention is an important mechanism for generating voluntary saccadic eye movements. Therefore, our result indicates that saccades are not only gaze but also an attention could be directed toward the experimenter side of the court even in the ball approach phase. This result supports our results about fixation and gaze targets as mentioned above.

On the other hand, our results have been contrary to previous research that saccade is directed ahead of the ball to predict the ball trajectories for intercept performance such as baseball (Kishita et al., 2020), cricket (Land and McLeod, 2000; Mann et al., 2013) and squash (Hayhoe et al., 2013). Therefore, our results would be the first to show a novel pattern of predictive visual patterns, that skilled table tennis players successfully execute constant forehand rallies without gazing at and paying attention to the entire ball trajectories.

## 5. Conclusion

This study examined predictive visual patterns of skilled table tennis players during constant forehand rallies. The results showed that participants tend to gaze at the ball when the experimenter hit the ball. We also found that the gaze target remained stationary on the experimenter side of the court even in the ball approach phase. Furthermore, saccades were directed toward the experimenter side of the court after fixation on the ball. These findings suggest that kinematic information about the opponent is important for successful forehand table tennis rallies. Taken together, skilled table tennis players are most likely to use unique visual patterns for interceptive sports players to estimate spatiotemporal information about the ball.

